# Activation of innate immunity by neutral polysaccharides from broccoli buds

**DOI:** 10.1101/2021.03.10.434731

**Authors:** Atsushi Miyashita, Keiko Kataoka, Kazuhisa Sekimizu

**Affiliations:** Institute of Medical Mycology, Teikyo University, Tokyo, Japan; Genome Pharmaceuticals Institute Co., Ltd., Tokyo, Japan

## Abstract

Edible substances that stimulate the innate immune system are good candidates for functional foods to improve human health. We have previously reported that acidic polysaccharides from broccoli extract exhibit immunostimulatory effects, but neutral polysaccharides have been overlooked. In the present study, we found that neutral polysaccharides have significantly stronger (higher specific activity) immunostimulatory activity than acidic polysaccharides. The hot water extract of broccoli showed the immunostimulatory activity in the silkworm muscle contraction assay, suggesting that it stimulates innate immunity *via* paralytic peptide pathway. The activity was concentrated in the buds, but not in the stems and stalk. The active substance was recovered in the flow-through fraction of diethylaminoethyl-cellulose column chromatography with neutral polysaccharides. The specific activity of the fraction was significantly higher than that of the acidic polysaccharides from broccoli reported previously. These results suggest that the neutral polysaccharide present in broccoli buds stimulates innate immunity and can be semi-purified by one-step chromatography.

## Introduction

Innate immunity is a mechanism that rapidly eliminates foreign or unwanted substances (bacteria, fungi, viruses, and cancer cells) from the body without the use of antibodies and is conserved in animals ranging from insects to mammals (1–5). We have reported that activation of innate immunity is accompanied by muscle contraction in the silkworm (6, 7). Furthermore, we have established a method to identify innate immunity activators in foods utilizing the silkworm muscle contraction as an easy bioassay system (8, 9). In addition, we have already reported that there are polysaccharides such as β-glucan that exhibit high specific activity using the silkworm muscle contraction assay system (9). These polysaccharides are expected to contribute to human health.

In our previous study, we found that the hot water extract of the edible part of broccoli showed higher specific activity in the muscle contractile system of the silkworm than other vegetables (10). In addition, the active component was ethanol precipitated, subjected to diethylaminoethyl (DEAE)-cellulose column chromatography, and purified by eluting the DEAE-bound fraction, revealing its structure to be a pectin-like acidic polysaccharide (10). On the other hand, some neutral polysaccharides in different plant materials have recently been reported to have physiological effects on the innate immune system (11–14), but there are no such reports for broccoli. Such neutral polysaccharides, if present in broccoli, should appear in the flow-through fraction of DEAE-cellulose column chromatography. In the present study, we examined whether there is bioactive neutral polysaccharide, which has not been reported so far, exists in the flow-through fraction of DEAE-cellulose column chromatography using the hot water extract of broccoli. In addition, in this study, we compared the innate immunity-stimulating activity of each edible part of broccoli in order to understand which part of broccoli contains the active substances.

## Materials and Methods

### Broccoli

Ten broccoli cultivars provided by Sakata Seed Corporation (Yokohama, Japan) were used in this study. For the cultivar No. 1, we performed two experiments using different batches of broccoli (one experiment for comparing three parts of broccoli, another for preparation of DEAE flow-through fraction). We have observed that different batches contain slightly different amount of the bioactivity (e.g., between harvest seasons; KS, pers. obs.)

### Hot water extract of broccoli polysaccharides from buds, stems, and stalks

Broccoli was divided into two edible parts (small flower buds, and stems) and an inedible part (stalks) with scissors, boiled in a warm bath for 2 minutes, and then immersed in cold water to cool well. The small flower buds were then separated from the stems using scissors, and the weight of buds, stems, and stalks were measured respectively. Fifteen grams of each part was taken and crushed in a polytron with 30 ml of pure water. Figure 1 is a photographic description of each part of the broccoli.

**Figure 1.**
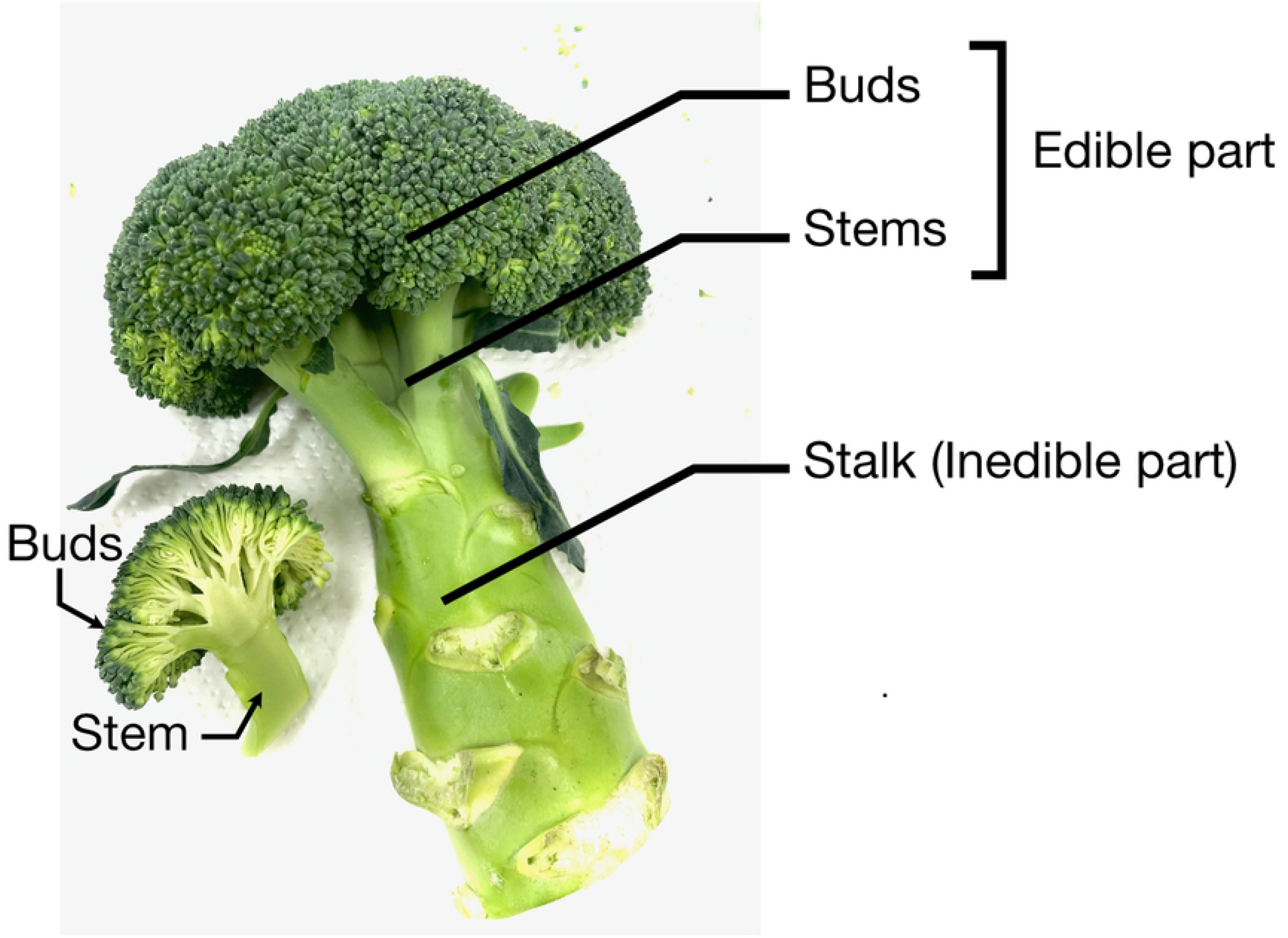
The buds, stems, and stalk of broccoli. The picture represents the three parts of broccoli (buds, stems, and stalks) as described in the Materials and Methods section. In this study, broccoli was broken down into these three parts and hot water extracts were prepared separately. The immunostimulatory activity of each part is summarized in Table 1.

**Table 1.** Total activity per 100 g wet weight of each part of broccoli. SD: Standard Deviation.

|  | Total activity per 100g (units) |  |  |
| --- | --- | --- | --- |
|  | buds | stems | stalk |
| No.1, replicate #1 | 1,200,000 | 43,000 | 7,100 |
| No.1, replicate #2 | 2,100,000 | 19,000 | 8,300 |
| No.1, replicate #3 | 1,700,000 | 58,000 | 4,300 |
| Mean<br>±SD | 1,600,000<br>±44,000<br>(n=3) | 98,000<br>±83,000<br>(n=3) | 6,600<br>±2,100<br>(n=3) |

### Silkworm muscle contraction assay

Muscle specimens were prepared by rearing 5^th^ instar silkworms on artificial diet for 5 days since the last ecdysis (10). The sample solution (50 μL) was injected into the muscle specimens using a tuberculin syringe (1mL) with a 27-gauge needle, and the body length was monitored for 10 min. The contraction value (C = (x-y)/x) was calculated for the length of the silkworm at the beginning of the experiment (x [cm]) and the length at the maximum contraction (y [cm]). The contraction values (C-values) for various sample volumes were plotted on a graph, and the sample volume with a C-value of 0.15 was estimated, and its activity was determined to be 1 unit.

### Preparation of flow-through fraction of DEAE-cellulose column chromatography

Please see Figure 2 for a simplified scheme of the whole process. Two times the amount of PCI (phenol: chloroform: isoamyl alcohol) was added to the hot water extract of broccoli (whole edible part), the aqueous layer was collected by centrifugation (8,000 rpm, 5 minutes), and two times the amount of ethanol was added to the aqueous layer fraction. The precipitate formed was collected by centrifugation (8,000 rpm, 10 min), dissolved in 10 mM Tris-HCl (pH 7.9) buffer, and applied to a DEAE-cellulose column (GE Healthcare, bed volume = 20 mL) equilibrated with the same buffer to collect the flow-through fraction (Figure 2).

**Figure 2.**
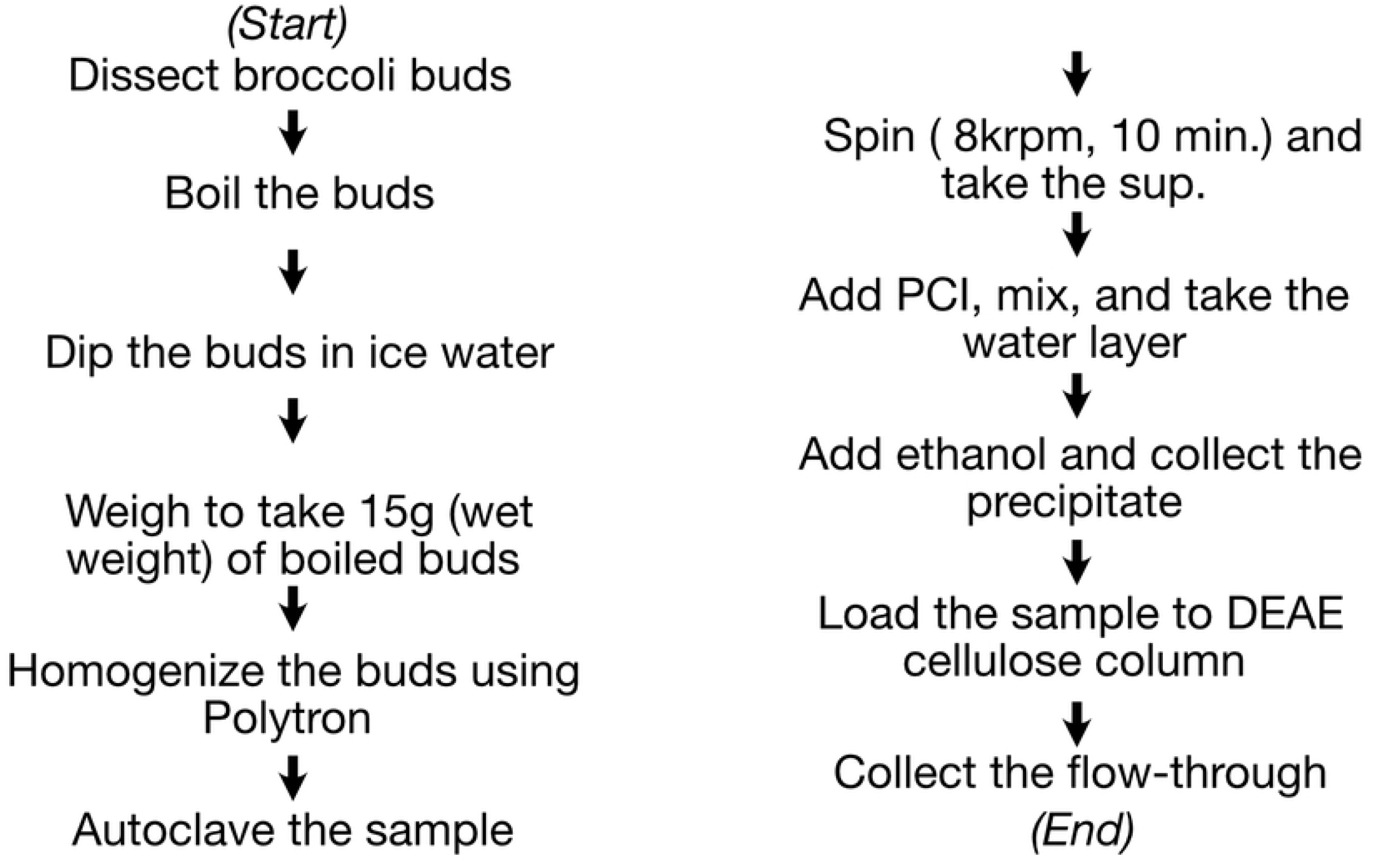
The scheme of the neutral polysaccharide preparation from broccoli. Shown in the panel is a simplified scheme for the preparation of neutral polysaccharides from broccoli: PCI, phenol-chloroform-isoamyl alcohol; DEAE, diethylaminoethyl. For a detailed description of each step, please refer to the Materials and Methods section.

## Results

### Innate immunity-stimulating substance is concentrated in the buds of broccoli

The total activity of each part of broccoli is shown in Table 1 (see Figure 1 for graphical details of broccoli parts). For every 100 g (wet weight) of start material, the hot-water extract of buds contained 1,600,000 ± 440,000 units (n=3), that of stems contained 98,000 ± 83,000 units (n=3), and that of stalk contained 6,600 ± 2,100 units (n=3). In percentage, these values represent 94 % for buds, 5.6 % for stems, and 0.4 % for stalk (see Table 1 for the results from three experimental replicates).

### Fractionation of hot water extracts of broccoli by DEAE cellulose column chromatography

The DEAE-cellulose flow-through fraction prepared from broccoli (cultivar No. 1) was determined for its immune-stimulating activity by the silkworm muscle contraction assay (Figure 3). Figure 3 demonstrates the results from three experimental replicates. We further tested several broccoli cultivars (Table 2), and all showed activity but at different specific activity value per sugar amount, ranging from a maximum of 13,000 units/mg sugar (cultivar No. 1) to a minimum of 840 units/mg sugar (cultivar No. 3).

**Figure 3.**
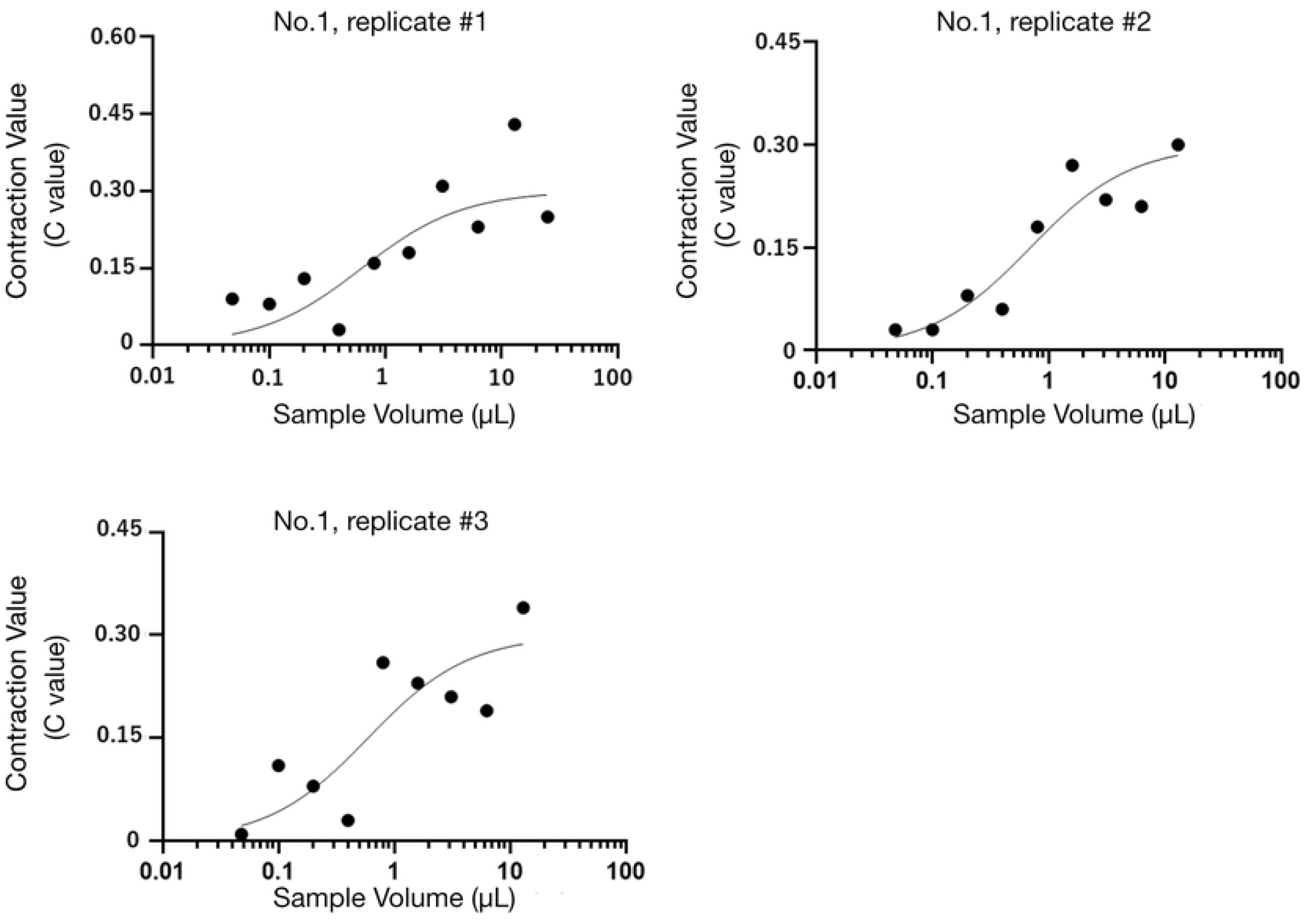
Immunostimulatory effect (silkworm muscle contraction assay) of the neutral polysaccharides of broccoli. The graph shows the dose-dependent immunostimulatory effect of a neutral polysaccharide sample prepared from broccoli cultivar No. 1 (see also Table 2 for a list of results for other cultivars). Three independent experimental replicates (replicate #1, replicate #2, and replicate #3) are shown in the panel. The vertical axis represents the C value (see Materials and Methods for how we determined the C values), and the horizontal axis represents the sample dose used in the assay.

**Table 2:** Innate immunity-stimulant activity of broccoli (ten cultivars)

| Cultivar # | Total activity per 100g of buds (units) | Sugar amount per 100g of buds (mg) | Specific Activity (units/mg sugar) |
| --- | --- | --- | --- |
| No.1 | 5,600,000 | 430 | 13,000 |
| No.2 | 1,400,000 | 400 | 3,600 |
| No.3 | 500,000 | 590 | 840 |
| No.4 | 390,000 | 150 | 2,600 |
| No.5 | 720,000 | 550 | 1,300 |
| No.6 | 3,400,000 | 600 | 5,700 |
| No.7 | 7,800,000 | 880 | 8,900 |
| No.8 | 16,000,000 | 1,800 | 8,700 |
| No.9 | 3,400,000 | 450 | 7,700 |
| No.10 | 1,000,000 | 360 | 2,900 |

## Discussion

In the present study, we examined the neutral polysaccharide fraction (the flow-through fraction of DEAE cellulose column chromatography using the hot water/PCI extract of broccoli as a start material) for its immunostimulatory activity. The results suggest the presence of neutral polysaccharides that act on innate immunity in the broccoli. In addition, in the present study, we compared the innate immunity-stimulating activity of three edible parts (buds, stems, and stalk) in order to understand the distribution of the active substance. As a result, it turned out that the buds contain most of the active molecules (94%, as demonstrated in Table 1), indicating the potential use of broccoli buds as ingredients for human health use. This is the first report for bioactive neutral polysaccharides in broccoli that stimulates innate immunity. Since the method presented in this study uses flow-through fractions, the sample preparation step is relatively simple. The advantage of using the flow-through fraction for sample preparation is that it does not require desalting (e.g., dialysis) as is the case when using the bound fraction of DEAE column chromatography. Nevertheless, further investigation is need for their chemical structures, and it remains to be seen what individual-level benefits (e.g., increase of immune gene expression, or acquisition of infection resistance) will bring to the actual infection process. Immune priming experiment using the silkworm (2, 5) is one of the possible approaches. Also, in the sample preparation step, the surface of broccoli was washed with boiling water before crushing, but because of the intricate structure of broccoli buds, it is not always possible to completely remove adhering matter (microorganisms, etc.). In this regard, it is necessary to purify the active substance and identify the chemical structure in the future studies.

## Acknowledgments

We thank Genome Pharmaceuticals Institute, Co. Ltd., for technical supports, and also thank Sakata Seed Corporation for providing materials. This work was supported by JSPS KAKENHI (Grant# 20K16253) to AM.

